# *Methanosarcina acetivorans* simultaneously produces molybdenum, vanadium, and iron-only nitrogenases in response to fixed nitrogen and molybdenum depletion

**DOI:** 10.1101/2021.06.03.447018

**Authors:** Melissa Chanderban, Christopher A. Hill, Ahmed E. Dhamad, Daniel J. Lessner

## Abstract

All nitrogen-fixing bacteria and archaea (diazotrophs) use molybdenum (Mo) nitrogenase to reduce dinitrogen (N_2_) to ammonia. Some diazotrophs also contain alternative nitrogenases that lack Mo: vanadium (V) and iron-only (Fe) nitrogenases. Among diazotrophs, the regulation and usage of the alternative nitrogenases in methanogens is largely unknown. *Methanosarcina acetivorans* contains *nif*, *vnf*, and *anf* gene clusters encoding putative Mo-, V-, and Fe-nitrogenases, respectively. This study investigated the effect of fixed nitrogen and Mo/V availability on nitrogenase expression and growth by *M. acetivorans*. The availability of Mo and V did not affect growth of *M. acetivorans* with fixed nitrogen but significantly affected growth with N_2_. *M. acetivorans* exhibited the fastest growth rate and highest cell yield during growth with N_2_ in medium containing Mo. Depletion of Mo (Fe-only condition) resulted in a significant decrease in growth rate and cell yield. The addition of V to Mo-depleted medium stimulated diazotrophic growth but was still less than growth in Mo-replete medium. qPCR analysis revealed transcription of the *nif* operon is only moderately affected by depletion of fixed nitrogen and Mo. However, *vnf* and *anf* transcription increased significantly when fixed nitrogen and Mo were depleted, with removal of Mo being the key factor. Immunoblot analysis revealed Mo-nitrogenase is produced when fixed nitrogen is depleted regardless of Mo availability, while V- and Fe-nitrogenases are produced only in the absence of fixed nitrogen and Mo. These results reveal that alternative nitrogenase production in *M. acetivorans* is tightly controlled and that all three nitrogenases can be simultaneously produced.

**IMPORTANCE:** Methanogens and closely related methanotrophs are the only archaea known or predicted to possess nitrogenase. As such, methanogens play critical roles in both the global biological nitrogen and carbon cycles. Moreover, methanogens are an ancient microbial lineage and nitrogenase likely originated in methanogens. An understanding of the usage and properties of nitrogenases in methanogens can provide new insight into the evolution of nitrogen fixation and aid in the development nitrogenase-based biotechnology. This study provides the first evidence that a methanogen can produce all three forms of nitrogenases, even simultaneously. Surprisingly, Mo-nitrogenase was produced in cells grown in the absence of Mo, indicating components of Mo-nitrogenase regulate or are needed to produce V- and Fe-nitrogenases in methanogens. The results provide a foundation to understanding the assembly, regulation, and activity of the alternative nitrogenases in methanogens.

## INTRODUCTION

Microbes are the primary drivers of the global biological nitrogen (N) cycle [1, 2]. For example, only select bacteria and archaea are capable of biological nitrogen fixation, whereby dinitrogen gas (N_2_) is reduced to ammonia (NH_3_), the preferred “fixed” form of N used directly by most organisms. The biological reduction of the triple bond of N_2_ is difficult and is catalyzed by nitrogenase, a unique metalloenzyme [3, 4]. To date, all known and predicted N_2_-fixing prokaryotes (diazotrophs) possess molybdenum (Mo) nitrogenase that contains a Mo atom within the unique iron (Fe) Mo-cofactor or M-cluster of the active site [5, 6]. Mo-nitrogenase consists of two components; the Fe protein, which contains a single iron-sulfur (Fe-S) cluster, and the MoFe protein that contains the active site FeMo-cofactor and the [8Fe-7S] P-cluster. The Fe protein, encoded by *nifH*, is the dinitrogenase reductase that donates electrons to the MoFe protein, the dinitrogenase composed of a heterotetramer of subunits encoded by *nifD* and *nifK*. Together NifH and NifDK catalyzes the energy intensive reduction of N_2_ as shown: N_2_ + 16ATP + 8e^−^ + 8H^+^ → 2NH_3_ + H_2_ + 16ADP + 16P_i_ [7]. As such, Mo-nitrogenase production and activity is highly regulated in diazotrophs and is only synthesized when a fixed N source is unavailable. When needed, Mo-nitrogenase is produced in high quantities and can comprise as much as 10% of the total protein of the cell [8].

In addition to having Mo-nitrogenase, some diazotrophs possess alternative nitrogenases that lack Mo [9, 10]. The vanadium (V) nitrogenase and the Fe-only (Fe) nitrogenase contain an active site FeV-cofactor and FeFe-cofactor, respectively, instead of FeMo-cofactor [11, 12]. The understanding of the genetic, biochemical, and catalytic properties of the alternative nitrogenases has primarily come from a few model bacteria (e.g., *Azotobacter vinelandii*). V-nitrogenase and Fe-nitrogenase have a similar subunit composition as Mo-nitrogenase, comprised of VnfH/VnfDK and AnfH/AnfDK subunits, respectively. However, a distinguishing feature of V- and Fe-nitrogenases is the presence of an additional subunit (G) that associates with the dinitrogenase component (i.e., VnfDGK and AnfDGK) [9, 11]. The precise role of the G subunit is unknown, but it is required for diazotrophy in the absence of Mo [13]. V- and Fe-nitrogenases are less efficient at reducing N_2_ than Mo-nitrogenase. More electron flux is directed to obligate H_2_ production during reduction of N_2_ by the alternative nitrogenases leading to substantially more ATP consumption. The V- and Fe-nitrogenases are estimated to consume 24 ATPs and 40 ATPs, respectively, during the reduction of a single N_2_ to 2NH_3_ [14, 15]. As such, alternative nitrogenases in bacteria are only produced when insufficient levels of Mo are present to support usage of Mo-nitrogenase. In studied bacteria that possess all three nitrogenases, the expression and activity of each nitrogenase is highly regulated in response to metal and fixed N availability [9, 16].

In addition to N_2_, nitrogenases from bacteria can reduce other double and triple-bonded substrates (e.g., CO, CO_2_, acetylene). Moreover, in the absence of another substrate, nitrogenase reduces protons to H_2_, a feature that has been exploited to use nitrogenase to produce H_2_ as a biofuel [17, 18]. The substrate, product, and activity profiles are also different between the three nitrogenases. The reduction of acetylene (C_2_H_2_) to ethylene (C_2_H_4_) is commonly used to measure nitrogenase activity [19]. Mo-nitrogenase reduces acetylene at a higher rate than both V- and Fe-nitrogenases, which also further reduce ethylene, producing ethane (C_2_H_6_) as a minor product [20]. Mo-nitrogenase does not produce ethane. Moreover, bacterial V-nitrogenase is more adept at reducing CO to alkanes, and the Fe-nitrogenase is better at reducing CO_2_ to CH_4_ [11, 21–23].

In contrast to bacterial diazotrophs, the regulation, assembly, and activity of nitrogenase, especially the alternative nitrogenases, is largely unknown in archaeal diazotrophs. Among archaea, only anaerobic methanogens and the closely related anerobic methanotrophs are known or predicted to fix N_2_ [5, 24, 25]. N_2_ fixation has been studied in a few species of methanogens. The primary models are the obligate CO_2_-reducing methanogen *Methanococcus maripaludis*, and the more versatile species *Methanosarcina mazei* and *Methanosarcina barkeri* [26, 27]. *Methanosarcina* species can grow using methylated compounds (e.g., methanol) and acetate, in addition to reducing CO_2_ with H_2_ [28]. *M. maripaludis* and *M. mazei* only contain Mo-nitrogenase, whereas strains of *M. barkeri* contain all three nitrogenases [29, 30]. Mo-dependent and V-dependent N_2_ fixation has been demonstrated in *M. barkeri* [31–33]. To our knowledge, diazotrophy under Fe-only conditions using the Fe-nitrogenase has not been documented for any methanogen. Previous research has primarily focused on elucidating the mechanisms that regulate the production and activity of Mo-nitrogenase in methanogens, revealing that the regulatory proteins used to control transcription and activity of Mo-nitrogenase are distinct from those used by most bacteria [34, 35]. Recently, small RNAs (sRNA) have also been demonstrated to play roles in N_2_ fixation and assimilation in methanogens [36, 37].

*Methanosarcina acetivorans* serves as an ideal model methanogen to understand the regulation and usage of the alternative nitrogenases in methanogens, since its genome encodes all three nitrogenases and it has a robust genetic system [38–41]. Recently, it was shown that *M. acetivorans* can fix N_2_ using Mo-nitrogenase. Like *M. maripaludis*, *M. mazei*, and *M. barkeri*, Mo-nitrogenase is only produced in *M. acetivorans* when cells are grown in the absence of a fixed N source (e.g., NH_4_Cl). Silencing of the *nif* operon in *M. acetivorans* using the recently developed CRISPRi-dCas9 system confirmed that Mo-nitrogenase is required for diazotrophy when cells are supplied Mo [41]. However, to our knowledge, the ability of *M. acetivorans* to fix N_2_ when Mo is not available has not been documented nor have the activities of *M. acetivorans* V-nitrogenase or Fe-nitrogenase been reported. Presumably, *M. acetivorans* produces V-nitrogenase and/or Fe-nitrogenase when both fixed N and Mo are limiting. An understanding of the properties of nitrogenases from methanogens could lead to new avenues for nitrogenase-based biofuel production and for the genetic engineering of crop plants capable of N_2_-fixation. In this study we show that *M. acetivorans* can grow by fixing N_2_ in the absence of Mo with production of both V- and Fe-nitrogenases. These results provide a foundation to understand the regulation and properties of the three nitrogenases in methanogens.

## RESULTS

### Organization of nitrogenase genes in *M. acetivorans* and prevalence of alternative nitrogenases in methanogens

The genome of *M. acetivorans* contains three separate nitrogenase gene clusters (**Fig. 1**), designated *nif*, *vnf*, and *anf*, encoding putative Mo-nitrogenase, V-nitrogenase, and Fe-nitrogenase, respectively. The gene arrangement of the *nif* cluster is similar to the characterized *nif* operons from *M. maripaludis*, *M. barkeri*, and *M. mazei* [30, 42, 43]. In addition to encoding the nitrogenase structural components (NifH and NifDK), the operon also encodes the regulatory proteins NifI_1_ and NifI_2_ and the FeMo-cofactor scaffold proteins NifEN [12, 44]. The *M. acetivorans vnf* cluster contains the same gene arrangement as *nif*, including its own regulatory and scaffold genes, but also includes *vnfG* and a homolog of *nifX*, designated *vnfX*. NifX is involved in FeMo-cofactor assembly in bacteria [12]. The gene arrangement of the *M. acetivorans anf* cluster is like the *vnf* cluster, except *anfH* encoding the putative Fe-protein is located divergent and downstream of *anfK*. The *anf* and *vnf* gene clusters are divergent in the chromosome of *M. acetivorans* (**Fig. 1**), indicating there could be coordinated regulation. Interestingly, the amino acid sequences of VnfH and AnfH are identical, indicating the same Fe-protein functions with both V- and Fe-nitrogenases. Also unique to the *anf* cluster is the presence of homologs of Anf3 and AnfO found in *anf* operons of bacteria. The precise functions of Anf3 and AnfO are unknown. Anf3 is essential for diazotrophy with the Fe-nitrogenase in *Rhodobacter capsulatus* [45]. An Anf3 homolog characterized in *A. vinelandii* is a heme- and FAD-binding oxidase that may protect the Fe-nitrogenase from oxygen [46].

**Figure 1.**
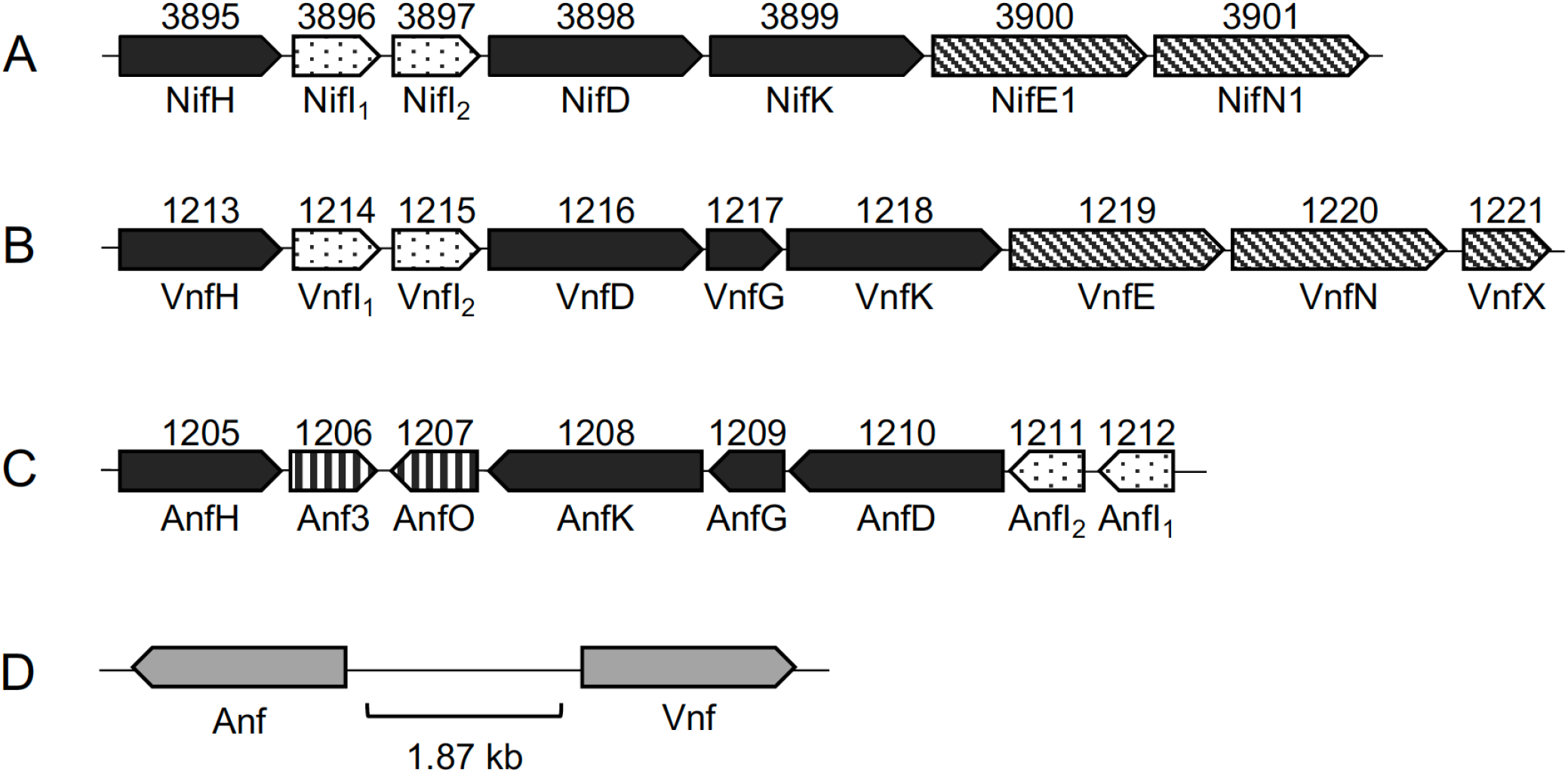
Arrangement of nitrogenase gene clusters in the genome of *M. acetivorans*. A) *nif*; Mo-nitrogenase, B) *vnf*; V-nitrogenase, C) *anf*; Fe-nitrogenase. The locus tag is above and the predicted protein below. Black arrows: nitrogenase subunits, diagonal striped arrows: cofactor assembly proteins, dotted arrows: regulatory proteins and vertical striped arrows: unknown function. D) the *vnf* and *anf* gene clusters are divergent in the chromosome as shown.

The *nif*, *vnf*, and *anf* gene clusters are widely distributed within genera of bacteria. However, nitrogenase genes are found only in a subset of archaea, restricted to methanogens and closely related anerobic methanotrophs. The *nif* operon is distributed across six of the seven orders of methanogens, whereas the *vnf* and *anf* genes are restricted to the Methanosarcinales, with few exceptions, namely *Methanobacterium lacus*, which contains a putative *anf* gene cluster [5, 24, 25]. Like bacteria, all methanogens that contain putative *vnf* and *anf* clusters also contain the *nif* operon. Of the 41 complete Methanosarcinales genome sequences currently available in the NCBI database, ~ 66 % contain the *nif* genes. Of those containing *nif*, ~ 44 % contain the *vnf* and/or *anf* genes (**Table 1**). The arrangement of the *vnf* and *anf* gene clusters are similar across the Methanosarcinales (**Fig. S1**). Of note is a hypothetical protein encoded by a gene between *vnfDGK* and *vnfEN* in several *Methanosarcina* species.

**Table 1.**
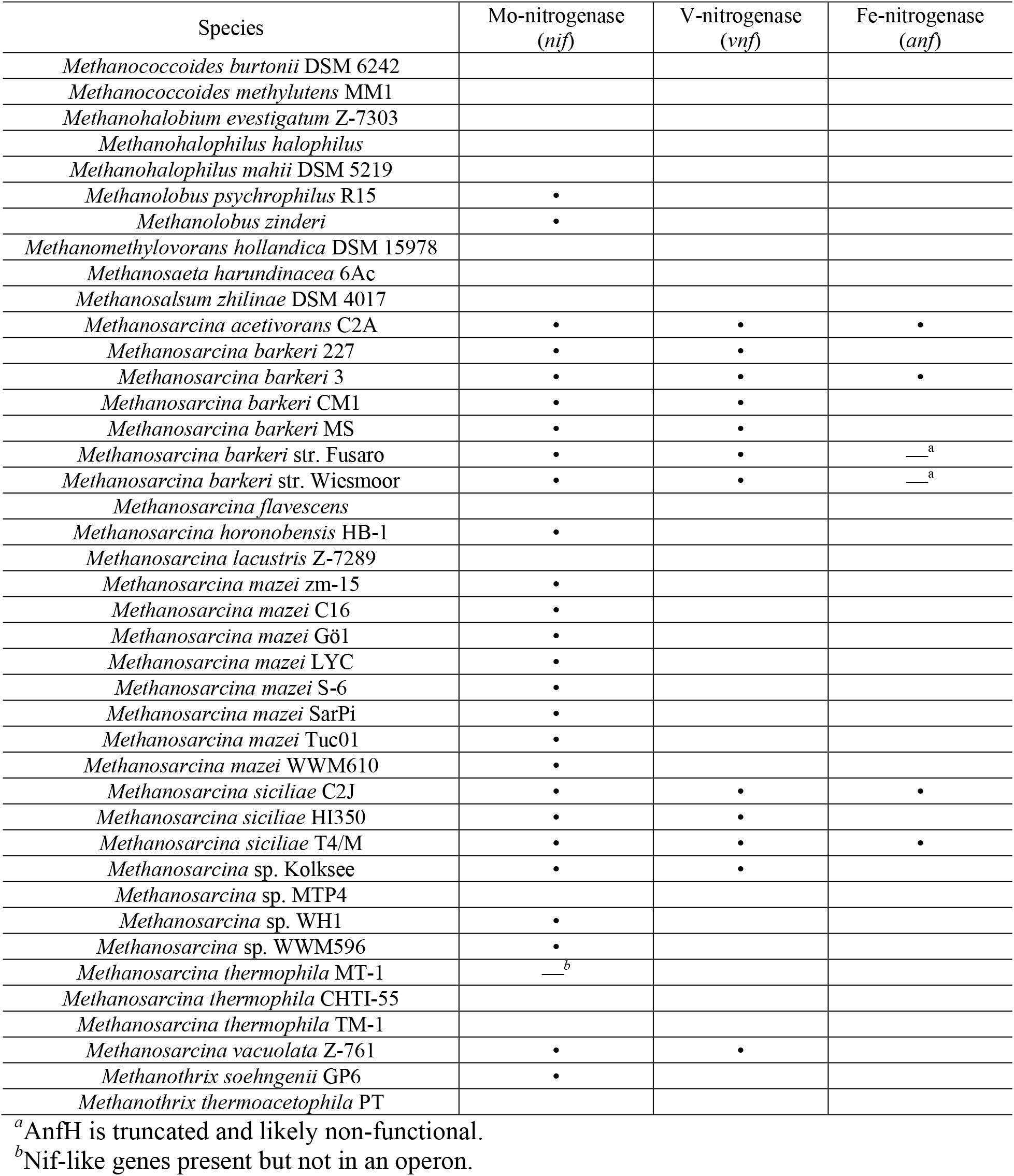
Nitrogenase distribution among genome-sequence Methanosarcinales.

### Molybdenum and vanadium availability affect diazotrophic growth of *M. acetivorans*

To ascertain the effect of molybdenum and vanadium availability on nitrogenase utilization by *M. acetivorans*, the pseudo-wild-type strain WWM73 (used for genetic analysis) [40] was passed in HS standard medium lacking Mo for >100 generations to deplete molybdate, the biological available form of Mo. Vanadium is not present in standard HS medium. Mo-deplete cells were used to inoculate Mo-deplete HS medium devoid of NH_4_Cl (fixed N source). Methanol was used as the carbon and energy source in all experiments. Molybdate, vanadate, and NH_4_Cl were added from sterile anaerobic stocks to separate cultures to compare the effect of Mo, V, and fixed N on growth and nitrogenase expression. Neither the depletion of Mo nor the addition of V affects the growth profile, generation time, or cell yield when NH_4_Cl is supplied as the fixed N source (**Fig. 2 and Table 2**). However, the depletion of Mo and the addition of V significantly affects the growth profile, generation time and cell yield in cultures without NH_4_Cl (diazotrophic). When *M. acetivorans* is provided Mo in the absence of NH_4_Cl, the generation time increases approximately 3-fold, and the cell yield decreases approximately 37% compared to non-diazotrophic cultures (**Table 2**). Diazotrophic cultures lacking Mo but provided V have an even longer generation time and further reduction in cell yield (~50% that of non-diazotrophic cultures). Diazotrophic growth is further impacted by the absence of both Mo and V, with an ~10-fold increase in generation time and an ~70 % reduction in cell yield compared to non-diazotrophic cultures (**Fig. 2 and Table 2**). Diazotrophic cultures lacking Mo also have an extended lag phase compared to diazotrophic cultures containing Mo (**Fig. 2 and Table 2**). These data reveal that *M. acetivorans* is capable of diazotrophy in the absence of Mo, and that V availability impacts N_2_ fixation. These results are consistent with *M. acetivorans* utilizing Mo-, V-, and Fe-nitrogenases to fix N_2_ according to Mo and V availability.

**Figure 2.**
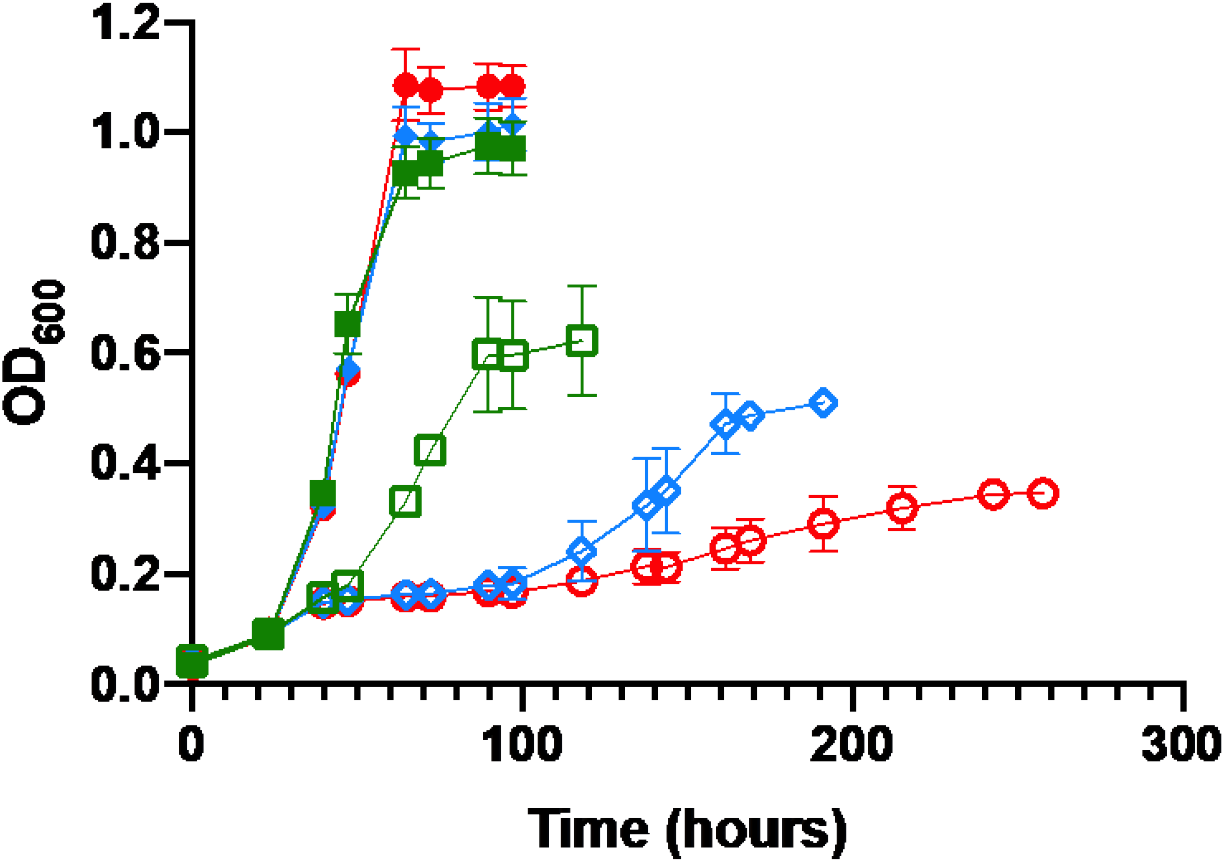
Comparison of the growth of *M. acetivorans* in the presence (closed) or absence (open) of NH_4_Cl in HS medium with Mo + Fe (green squares), V + Fe (blue diamonds), or Fe alone (red circles). Error bars represent mean ± 1 SD from at least three biological replicates.

**Table 2.** Effect of metal and NH_4_Cl availability on growth of *M. acetivorans* with methanol.

| Relevant Metals | Nitrogen Source | Lag time<sup>a</sup><br>(hours) | Generation Time<sup>b</sup><br>(hours) | Cell Yield<sup>b</sup><br>(cells/mL) |
| --- | --- | --- | --- | --- |
| Mo + Fe | NH<sub>4</sub>Cl | 30 | 8.2 ± 0.5 | 3.02 x 10<sup>8</sup> |
|  | N<sub>2</sub> | 48 | 28.5 ± 4 | 1.92 x 10<sup>8</sup> |
| V + Fe | NH<sub>4</sub>Cl | 30 | 8.5 ± 0.1 | 3.08 x 10<sup>8</sup> |
|  | N<sub>2</sub> | 90 | 44.5 ± 4.1 | 1.53 x 10<sup>8</sup> |
| Fe only | NH<sub>4</sub>Cl | 30 | 8.7 ± 0.1 | 3.34 x 10<sup>8</sup> |
|  | N<sub>2</sub> | 96 | 82 ± 4.1 | 9.88 x 10<sup>7</sup> |

<sup>a</sup>Approximate time until the first observed increase in OD<sub>600</sub>.
<sup>b</sup>Generation time and cell yield represent the mean ± 1 SD from at least three biological replicates.

### Methylotrophic methanogenesis is not altered by diazotrophy or the availability of molybdenum or vanadium

Growth of *M. acetivorans* with methanol utilizes the methylotrophic pathway of methanogenesis, where one methyl group of methanol is oxidized to CO_2_, and the resulting three electron pairs are used to reduce three additional methyl groups to CH_4_ [47]. To determine if diazotrophy and metal availability affect the flux of carbon during methylotrophic methanogenesis, contributing to the slower growth rate and lower cell yields in the absence of Mo, total CH_4_ was determined after the cessation of growth of non-diazotrophic and diazotrophic cultures. Similar amounts of CH_4_ were observed across all growth conditions (**Table 3**), revealing N_2_ fixation and differences in Mo and V availability does not significantly alter the flux of carbon during methylotrophic methanogenesis. Therefore, the observed hierarchical decrease in cell yields during diazotrophic growth under Mo + Fe, V + Fe, or Fe-only conditions (**Table 2**) is not due to decreased energy availability from altered methanogenesis but is likely due to the increased ATP consumption needed to support N_2_ reduction by Mo-, V-, and Fe-nitrogenases, as seen in bacteria [9].

**Table 3.**
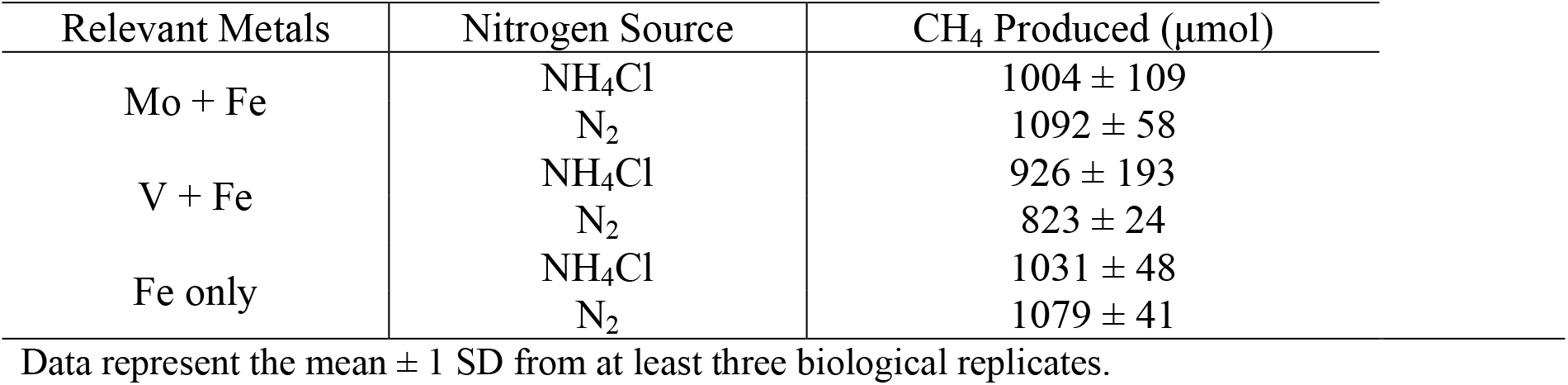
Effect of metal and NH_4_Cl availability on total CH_4_ production by *M. acetivorans* with methanol.

| Relevant Metals | Nitrogen Source | CH<sub>4</sub> Produced (μmol) |
| --- | --- | --- |
| Mo + Fe | NH<sub>4</sub>Cl | 1004 ± 109 |
|  | N<sub>2</sub> | 1092 ± 58 |
| V + Fe | NH<sub>4</sub>Cl | 926 ± 193 |
|  | N<sub>2</sub> | 823 ± 24 |
| Fe only | NH<sub>4</sub>Cl | 1031 ± 48 |
|  | N<sub>2</sub> | 1079 ± 41 |

Data represent the mean ± 1 SD from at least three biological replicates.

### Molybdenum availability affects the expression of V-nitrogenase and Fe-nitrogenase but not Mo-nitrogenase in *M. acetivorans*

Previous results demonstrated that Mo-nitrogenase is not produced in *M. acetivorans* cells grown in the presence of NH_4_Cl. Removal of NH_4_Cl results in a modest increase in *nif* transcription and production of Mo-nitrogenase, allowing growth with N_2_. Repression of the *nif* operon by dCas9 abolished the ability to grow with N_2_ in medium containing Mo [41]. To determine the effect of fixed N and Mo depletion on Mo-nitrogenase, V-nitrogenase and Fe-nitrogenase expression, qPCR was performed using primers specific for *nifD*, *vnfD*, and *anfD* (**Table S1**) to analyze transcript abundance in cells grown in medium with or without NH_4_Cl and containing Mo + Fe, V + Fe, or Fe only (**Fig. 3**). An increase in transcript abundance for *nifD* and *vnfD* was observed in cells grown in Mo + Fe medium without NH_4_Cl, relative to the transcript abundance in cells grown with NH_4_Cl (**Fig. 3A**). However, only the fold change for *vnfD* was significant. Comparison of *nifD*, *vnfD*, *anfD* transcript abundance from cells grown with V + Fe showed a significant fold change for *vnfD* and *anfD* (**Fig. 3B**). The transcript abundance of *vnfD* is ~180-fold higher in cells grown in V + Fe medium without NH_4_Cl compared to cells grown with NH_4_Cl. Transcript abundance for *anfD* is ~60-fold higher in cells grown in V + Fe medium without NH_4_Cl compared to cells grown with NH_4_Cl. In contrast, only a slight increase (~3-fold) was observed for *nifD* transcript abundance. Like the transcript abundance of *vnfD* and *anfD* in cells grown with V + Fe, cells grown in Fe-only medium lacking NH_4_Cl had a significant increase in *vnfD* and *anfD* transcript abundance compared to cells grown with NH_4_Cl (**Fig. 3C)**. No change in the expression of *nifD* was detected in cells grown in Fe-only medium lacking NH_4_Cl relative to that with NH_4_Cl (**Fig. 3C**).

**Figure 3.**
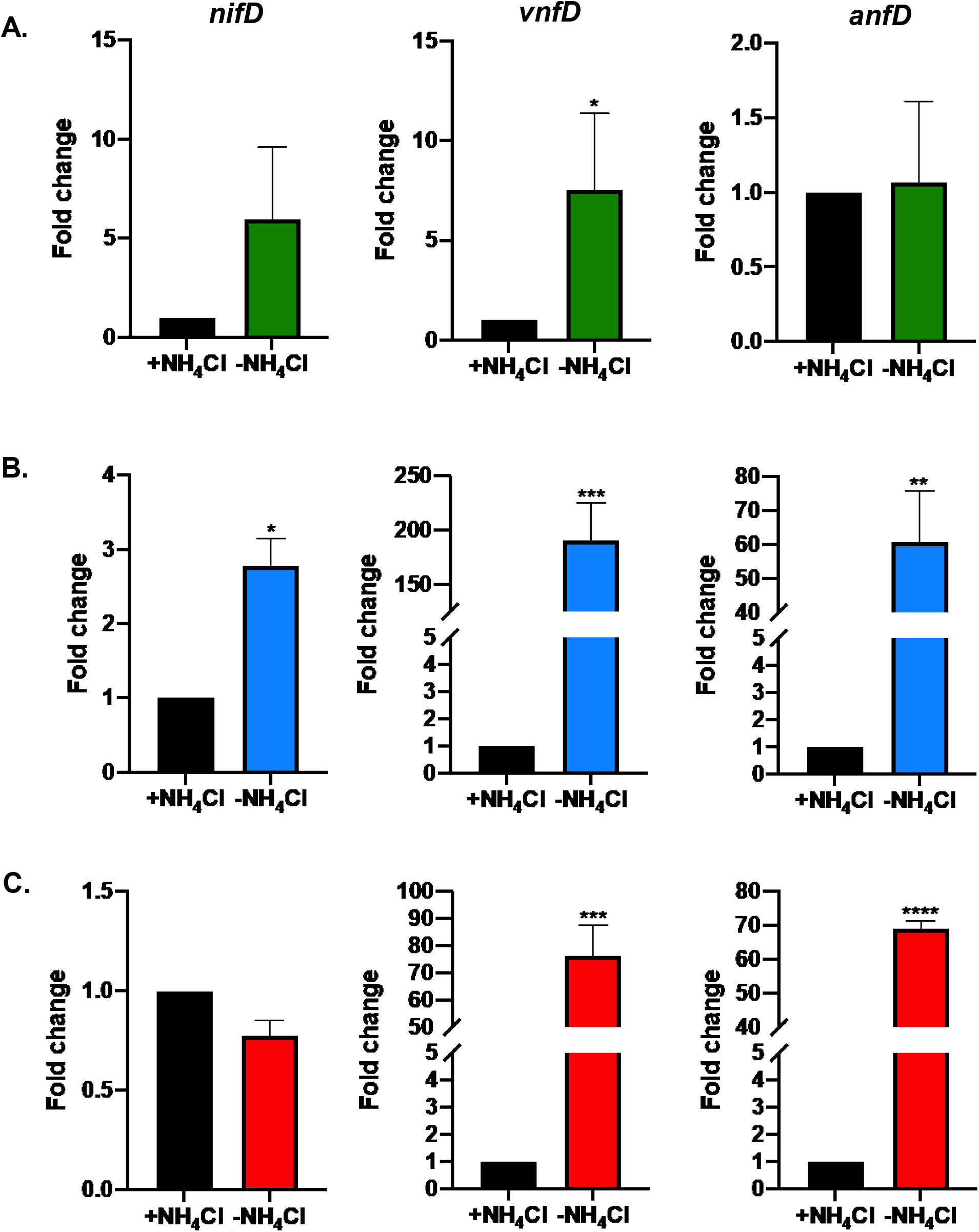
Effect of fixed N availability on the transcription of the *nif*, *vnf* and *anf* gene clusters in *M. acetivorans* as determined by qPCR. The relative abundance of *nifD*, *vnfD*, and *anfD* transcripts in *M. acetivorans* cells grown with NH_4_Cl (normalized to one) were compared to cells grown without NH_4_Cl. *M. acetivorans* was grown with methanol in HS medium containing A) Mo + Fe B) V + Fe or C) Fe only. Error bars represent mean ± 1 SD for two technical replicates and three biological replicates. *, P < 0.05; **, P < 0.01; ***, P < 0.001; ****, P < 0.0001.

To further determine the effect of Mo removal on transcription of each nitrogenase gene cluster, the fold change in *nifD*, *vnfD* and *anfD* transcript abundance was also calculated by comparing the relative abundance in cells grown in V + Fe or Fe-only medium to the transcript abundance in cells grown in Mo + Fe medium (**Fig. 4**). The expression of *nifD* did not significantly change in cells grown in medium with or without Mo, regardless of the presence or absence of NH_4_Cl (**Fig. 4A**). However, removal of Mo significantly affected the transcription of both *vnfD* and *anfD* in cells grown with or without NH_4_Cl (**Fig. 4B-C**). The transcript abundance of *vnfD* is highest in cells grown in Fe-only medium, with the fold-change higher than when V is present. A similar pattern was observed for the expression of *anfD*. However, the fold change in expression of *anfD* in cells grown with Fe only compared to Mo + Fe was much higher (~300-600-fold). These results indicate there is significant regulatory control of transcription of the *vnf* and *anf* gene clusters, whereas there is only modest transcriptional control of the *nif* operon. The results also show that the depletion of Mo is the key signal that increases transcription of the *vnf* and *anf* gene clusters. Removal of a fixed N source (NH_4_Cl) when Mo is available has only a slight effect on the transcription of the *vnf* and *anf* gene clusters (**Fig. 3A**).

**Figure 4.**
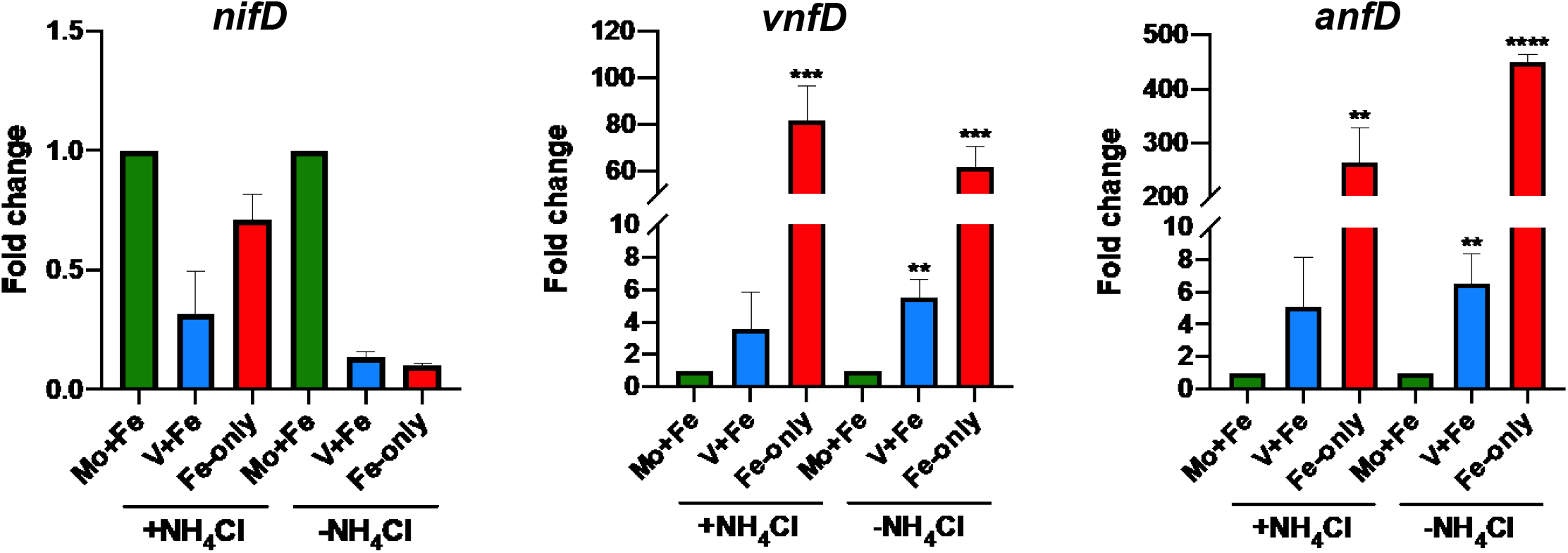
Effect of molybdenum availability on the transcription of the *nif*, *vnf* and *anf* gene clusters in *M. acetivorans* as determined by qPCR. The relative abundance of A) *nifD*, B) *vnfD*, and C) *anfD* transcripts in cells grown with molybdenum (normalized to one) were compared to cells grown without molybdenum. Error bars represent mean ± 1 SD for two technical replicates and three biological replicates. *, P < 0.05; **, P < 0.01; ***, P < 0.001; ****, P < 0.0001.

The production of Mo-, V-, and Fe-nitrogenases in *M. acetivorans* grown under the same conditions for qPCR analysis was determined by Western blot using antibodies specific to NifD, VnfD, and AnfD (**Fig. 5**). Consistent with previous results [41], NifD was only detected in lysate from *M. acetivorans* cells grown in Mo + Fe medium lacking NH_4_Cl. Neither VnfD nor AnfD were detected in lysate from cells grown in Mo + Fe medium regardless of the presence or absence of NH_4_Cl. However, both VnfD and AnfD were detected in lysate from cells grown in Mo-depleted medium lacking NH_4_Cl. Interestingly, NifD was also detected in lysate from cells grown in Mo-deplete medium. The availability of V does not appear to affect production of VnfD or AnfD. These results indicate that both the depletion of fixed N and Mo are required for production of V-nitrogenase and Fe-nitrogenase in *M. acetivorans*.

**Figure 5.**
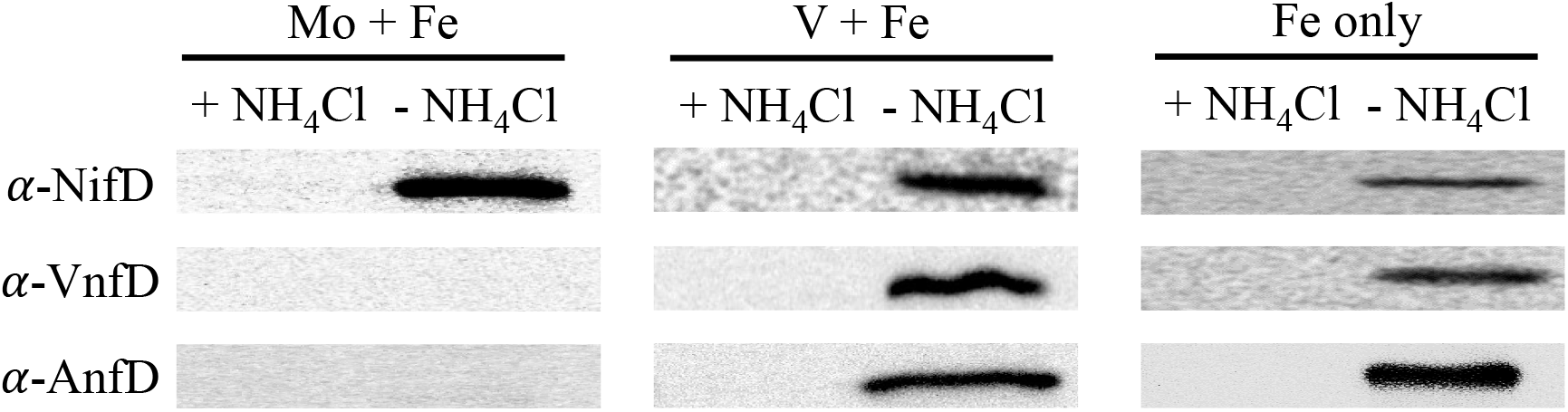
Western blot analysis using NifD-, VnfD-, and AnfD-specific antibodies on lysate from *M. acetivorans* cells grown with or without NH_4_Cl and the indicated metals.

## DISCUSSION

The regulation, assembly, and activity of the three forms of nitrogenase is well understood in diazotrophic bacteria, especially in the principal model *A. vinelandii* that contains all three nitrogenases. *A. vinelandii* is an obligate aerobe; thus, in addition to nitrogenase structural proteins, *A. vinelandii* requires accessory proteins to prevent oxidative damage to nitrogenase and to integrate nitrogen fixation into central metabolism. At least 82 genes are predicted to be involved in the formation and regulation of Mo-, V-, and Fe-nitrogenases in *A. vinelandii* [16]. Moreover, there is complex regulatory control over hierarchal nitrogenase expression, with only one nitrogenase produced at a time. When fixed N is absent and Mo is available, Mo-nitrogenase is preferentially produced over V- and Fe-nitrogenase, followed by V-nitrogenase if Mo is absent and V is present. If neither Mo nor V is available, then Fe-nitrogenase is produced [24]. Among methanogens, the alternative nitrogenases are restricted primarily to the Methanosarcinales, the most metabolically diverse methanogens with the largest genomes. Nonetheless, the genomes of sequenced Methanosarcinales contain simpler nitrogenase gene clusters and lack many of the accessory and regulatory proteins found in *A. vinelandii* and other diazotrophic bacteria [25]. The formation and regulation of the alternative nitrogenases is likely simpler in methanogens compared to aerobic diazotrophic bacteria. The results presented here demonstrate that *M. acetivorans* produces all three nitrogenases and is capable of diazotrophy in the absence of available Mo and V (Fe-only condition). To our knowledge, this is first direct evidence of a methanogen producing an Fe-nitrogenase and capable of diazotrophy in the absence of Mo or V.

Like other diazotrophs, *M. acetivorans* only produces nitrogenase in the absence of fixed N. The diazotrophic growth profiles of *M. acetivorans* correlate with reported ATP requirements by Mo-, V-, and Fe-nitrogenase from bacteria [14]. *M. acetivorans* has the fastest growth rate and highest cell yield during diazotrophic growth when utilizing only Mo-nitrogenase. Only a modest increase in transcription of the *nif* operon was observed in response to fixed N depletion. The high basal level of transcription of the *nif* operon likely allows *M. acetivorans* to be poised for rapid Mo-nitrogenase production. The relatively short lag time before the onset of diazotrophic growth in Mo + Fe medium (**Table 2** and **Fig. 2**) supports the rapid production of Mo-nitrogenase.

The results indicating minimal transcriptional control of the *nif* operon further support that post-transcriptional regulation is a key factor controlling Mo-nitrogenase production. Previous studies investigated the role of NrpR in regulating the expression of Mo-nitrogenase in *M. acetivorans*. NrpR is the repressor of the *nif* operon in methanogens and indirectly senses fixed N availability by directly sensing intracellular 2-oxogluatrate levels [48]. A mutant strain of *M. acetivorans* where *nrpR* transcription was silenced using the CRISPRi-dCas9 system revealed that the depletion of NrpR results in an increase in the transcription of the *nif* operon, but the mutant still fails to produce detectable nitrogenase when grown with fixed N [41]. In *Methanosarcina mazei*, a small RNA (sRNA_154_) is exclusively expressed when fixed N is limiting and functions to stabilize the polycistronic mRNA produced from the *nif* operon [36]. The genome of *M. acetivorans* encodes a sRNA_154_ homolog, indicating similar post-transcriptional regulation of the *nif* operon. Interestingly, removal of Mo did not significantly alter transcription of the *nif* operon or the production of nitrogenase (**Fig. 4A and 5**). Therefore, the critical and likely only signal for Mo-nitrogenase production in *M. acetivorans* is fixed N limitation. This is distinct from diazotrophic bacteria that contain V- and Fe-nitrogenases. For example, *A. vinelandii* and the purple non-sulfur phototroph *Rhodopseudomonas palustris* both stop producing Mo-nitrogenase when Mo is depleted [24, 49].

While Mo-depletion had little effect on Mo-nitrogenase expression, it is critical for the expression of V- and Fe-nitrogenase in *M. acetivorans*. Both fixed N and Mo depletion are required for production of V-nitrogenase and Fe-nitrogenase (**Fig. 5**). Importantly, Mo depletion resulted in a significant increase in the relative transcript abundance of *vnfD* and *anfD* (**Fig. 3 and 4).** Thus, unlike production of Mo-nitrogenase, transcriptional regulation is a key mechanism to control production of V- and Fe-nitrogenases in *M. acetivorans*. The overall transcript abundance profiles for *vnfD* and *anfD* are similar across all growth conditions. Mo depletion appears to be a key effector as cells grown with NH_4_Cl exhibited a significant increase in transcript abundance of *vnfD* and *anfD* (**Fig. 4**). Nonetheless, neither VnfD nor AnfD were detected in cells grown with NH_4_Cl in Mo-depleted medium (**Fig. 5**), indicating post-transcriptional regulation of *vnf* and *anf* genes is also likely involved. Unexpectedly, in the absence of Mo, the presence of V does not increase the transcript abundance of *vnfD* and *anfD* as much as the increase during Fe-only conditions (**Fig. 4**). The role V plays in nitrogenase regulation is unknown in most diazotrophs. Nevertheless, when comparing the effect of fixed N depletion, a large relative fold change in transcript abundance for *vnfD* and *anfD* was observed in cells grown in V + Fe medium (**Fig. 3B**). Expression of the *vnf* and *anf* operons in *A. vinelandii* in the absence of Mo results in the production of either V-nitrogenase or Fe-nitrogenase depending on V availability, but not both. In contrast, V availability had no effect on V-nitrogenase or Fe-nitrogenase production in *M. acetivorans*, as each was produced in cells grown in Mo-depleted medium (**Fig. 5**). Notably, VnfH and AnfH are identical in amino acid sequence, indicating a single dinitrogenase reductase (VnfH/AnfH) can support the *in vivo* activities of separate dinitrogenases (VnfDGK and AnfDGK). While the expression results cannot distinguish which nitrogenase is active/functional, the growth profiles are consistent with the more-efficient V-nitrogenase active in cells grown in V + Fe medium and the less-efficient Fe-nitrogenase active in cells grown in Fe-only medium (**Fig. 2**).

Production of both V-nitrogenase and Fe-nitrogenase in *M. acetivorans* clearly requires fixed N depletion since neither VnfD nor AnfD were detected by immunoblot in lysate from cells grown with NH_4_Cl regardless of Mo availability. Regulation of V-nitrogenase and Fe-nitrogenase expression in response to fixed N availability does not likely involve direct control of *vnf* and *anf* transcription since fixed N depletion in the presence of Mo did not alter *anfD* transcript abundance and only had a modest effect on *vnfD* transcript abundance (**Fig. 3A**). These results are consistent with the promotor regions of both the *vnf* and *anf* gene clusters lacking the identified NrpR operator sequence [50]. The promoter regions also lack identified binding sites for NrpA, an activator of the *nif* operon in *M. mazei*, for which *M. acetivorans* encodes two homologs (MA0545 and MA0546) [51]. Thus, post-transcriptional regulation is likely the primary mechanism of control of V-nitrogenase and Fe-nitrogenase production in response to fixed N availability. It is possible sRNA_154_, or another sRNA, is responsive to fixed N depletion and functions to stabilize *vnf* and *anf* mRNAs, which allows for V-nitrogenase and Fe-nitrogenase production only when fixed N is depleted.

Mo availability is the key factor controlling transcription of both the *vnf* and *anf* gene clusters in *M. acetivorans*. In non-diazotrophic (e.g., *E. coli*) and diazotrophic bacteria, the molybdate-responsive transcriptional regulator ModE controls the expression of the high-affinity molybdate transporter ModABC as well as Mo-dependent enzymes [52]. In *A. vinelandii*, ModE indirectly represses expression of both V-nitrogenase and Fe-nitrogenase by directly repressing the transcription of the genes encoding the regulators VnfA and AnfA. VnfA activates transcription of the *vnf* operon and AnfA activates transcription of the *anf* operon in *A. vinelandii* [52]. The genome of *M. acetivorans* encodes several homologs of ModABC (MA0325-27, MA1235-37, and MA2280-82), including additional homologs of ModBC (MA3902-03) downstream of the *nif* operon. *M. acetivorans* contains a ModE homolog (MA0283) but lacks homologs to VnfA and AnfA. Potential ModE-binding sites are located upstream of *vnfH* and *anfI_1_*, the first genes in the *vnf* and *anf* gene clusters [53]. Therefore, it is highly plausible that ModE is responsible for repressing transcription of *vnf* and *anf* when sufficient Mo is available to support Mo-nitrogenase activity. Depletion of Mo (corepressor) likely results in removal of DNA-bound ModE and de-repression of transcription of the *vnf* and *anf* gene clusters, leading to the simultaneous production of V-nitrogenase and Fe-nitrogenase in *M. acetivorans*. The results are consistent with this regulatory mechanism. Interestingly, the starter inoculum used in all expression studies was maintained in Mo-deplete medium, which should result in an increase in *vnf* and *anf* transcription even during growth with NH_4_Cl (**Fig. 4**). As such, the starter inoculum should be primed to use the alternative nitrogenases once fixed N is depleted, yet there was a much longer lag period before the onset of growth in Mo-deplete medium compared to the onset of growth in Mo-deplete medium with added Mo (**Table 2 and Fig. 2**). This result indicates that there are likely other unknown regulatory factors involved in controlling the production of V-nitrogenase and Fe-nitrogenase in response to fixed N and Mo depletion.

The simultaneous production of all three nitrogenases in *M. acetivorans* during diazotrophy in Mo-deplete medium raises interesting questions. Why would *M. acetivorans* continue to produce Mo-nitrogenase under conditions when the enzyme is likely not functional? One plausible explanation is that because the energy conservation (i.e., ATP generation) during methanogenesis by *M. acetivorans* is significantly lower even during optimal conditions compared to studied diazotrophic bacteria [54], that *M. acetivorans* continues to produce Mo-nitrogenase when fixed N is limiting regardless of Mo availability to be poised to use the most efficient nitrogenase. However, we cannot rule out that a small amount of residual Mo is present in the Mo-deplete medium that maintains expression of Mo-nitrogenase. But it is unlikely that this is the case since both V-nitrogenase and Fe-nitrogenase are produced in Mo-deplete medium, indicating Mo removal is sufficient to induce expression of the less efficient nitrogenases. Moreover, *M. acetivorans* failed to grow for more than one day in Mo-deplete medium after residual fixed N was depleted, consistent with insufficient Mo to support Mo-nitrogenase activity.

Another plausible explanation for the continued production of Mo-nitrogenase in Mo-deplete medium is that Mo-nitrogenase proteins are required for the formation of functional V-nitrogenase and Fe-nitrogenase. NifH, in addition to providing electrons to NifDK during N_2_ reduction, serves multiple roles in nitrogenase maturation in bacteria. For example, NifH is involved in the synthesis of the complex metalloclusters within NifDK (e.g., P-cluster) [3, 12, 55]. Therefore, NifH could be required for metallocluster synthesis in VnfDGK and AnfDGK. Although VnfEN scaffold proteins are encoded in the *vnf* gene cluster, it is also possible NifEN is needed for metallocluster synthesis in VnfDGK and/or AnfDGK. Alternatively, inactive NifDK may serve a regulatory role in controlling the production of active V-nitrogenase and Fe-nitrogenase.

Finally, the simultaneous production of all three nitrogenases under Mo-deplete conditions begs the question, which nitrogenase(s) are functional? Although only NifD, VnfD, and AnfD were detected in cells growing in Mo-deplete medium, it is likely that NifDK, VnfDGK, and AnfDGK complexes are present since NifD is unstable in the absence of NifK [56]. Therefore, metal-dependent regulation of metallocluster insertion into NifDK, VnfDGK, and AnfDGK may control which nitrogenase is active. NifDK likely lacks FeMo-cofactor when produced in cells growing in Mo-deplete medium, while VnfDGK likely lacks FeV-cofactor when produced in the absence of V. AnfDGK could contain the FeFe-cofactor cluster regardless of the presence of V and always be active in cells grown in Mo-deplete medium. Moreover, the formation of hybrid nitrogenases is possible, as both VnfDGK and AnfDGK can incorporate the FeMo-cofactor resulting in a functional hybrid nitrogenase [57, 58]. It is unlikely that NifDK can incorporate the FeV-cofactor or FeFe-cofactor, although this cannot be ruled out. Importantly, mutant analysis using the CRISPR-Cas9 and CRISPRi-dCas9 systems [39, 41] can help address many of these questions. Overall, the results from this study highlight the utility of *M. acetivorans* as a model to understand the regulation, maturation, and activity of the three forms of nitrogenase in methanogens.

## Acknowledgments

We thank Tom Deere for helpful discussions and assistance with gas chromatography. This work was supported in part by DOE Biosciences grant number DE-SC0019226 (DJL), NSF grant number MCB1817819 (DJL), NSF Graduate Research Fellowship under grant number 1842401 (MC), and the Arkansas Biosciences Institute (DJL), the major research component of the Arkansas Tobacco Settlement Proceeds Act of 2000.

## MATERIALS AND METHODS

### *M. acetivorans* strains and growth

*M. acetivorans* strain WWM73, a pseudo-wild type strain used for genetic manipulation [40], was used for all experiments. Anoxic high-salt (HS) medium was prepared as previously described with some modifications [59]. To prepare Mo-deplete HS medium, all glassware was washed twice with 1 M HCl, once with 1 M H_2_SO_4_, and then rinsed with ultrapure water to remove any residual molybdate prior to use. NH_4_Cl and molybdate were omitted and the HS medium was reduced with 1.5 mM DTT. Methanol, NH_4_Cl, sodium sulfide (Na_2_S), sodium molybdate (Na_2_MoO_4_), and sodium vanadate (Na_3_VO_4_) were added from anoxic sterile stocks using sterile syringes prior to inoculation. *M. acetivorans* strain WWM73 was grown in Balch tubes containing 10 ml of HS medium with 125 mM methanol and 0.025 % Na_2_S (w/v). Molybdate (1 μM), vanadate (1 μM), and NH_4_Cl (18 mM) were added to cultures as indicated. *M. acetivorans* strain WWM73 was grown for more than 100 generations in Mo-depleted HS medium containing methanol and NH_4_Cl prior to the growth experiments. Growth was measured by monitoring optical density at 600 nm (OD_600_) using a spectrophotometer. Cell density was determined from OD_600_ using a standard curve generated by direct cell counts with a hemocytometer.

### Quantitative PCR analysis of gene expression

*M. acetivorans* cells were harvested during mid-log phase (0.3-0.4 OD_600_) by anaerobic centrifugation of 4-8 mL of culture. Cell pellets were resuspended in 1 mL Trizol and frozen at −80 °C. RNA was extracted using the Zymo Direct-zol Miniprep kit (#R2052) and further purified using the Invitrogen DNA-free DNA Removal Kit (#AM1906). cDNA was generated using the Bio-Rad iScript Select cDNA Synthesis Kit (#1708896). qPCR primers were designed using Geneious Prime (Supplemental Table 1). qPCR of three biological replicates and two technical replicates was performed with the SsoAdvanced Universal SYBR Green Supermix (Bio-Rad, #1725271). Relative quantification was determined using the 2^−ΔΔCq^ method.

### Western blot analysis

Separate custom polyclonal antibodies specific for *M. acetivorans* NifD, VnfD, or AnfD were generated using the PolyExpress Silver package (two epitopes) from Genscript. Specificity of the antibodies was confirmed using recombinant NifD, VnfD, and AnfD expressed in *E. coli* (data not shown). *M. acetivorans* cells were harvested during mid-log phase (0.3-0.4 OD_600_) by aerobic centrifugation (8500 x *g* for 10 minutes at 4°C) of 6 mL of culture. The cell pellet was resuspended in 50 mM Tris, 150 mM NaCl pH 7.2 with 1 mM PMSF and 1 mM benzamidine, normalized based on OD_600_, and frozen at −80°C. Whole cell lysate was generated by five freeze/thaw cycles and a one hour DNase (5 μg) treatment at 37°C. Protein concentration was determined using the Bradford assay. After blocking for one hour in TBST (20 mM Tris, 150 mM NaCl, 0.1% Tween pH 7.6) with 5% milk, membranes were incubated for 18 hours with the primary antibodies specific for NifD, VnfD, or AnfD, then washed three times with TBST. Membranes were then incubated with an HRP-conjugated secondary antibody (Promega) for one hour, followed by three washes with TBST. Finally, membranes were visualized using an enhanced chemiluminescent reagent (Thermo Scientific) and an Alpha Innotech imaging system.

### Methane determination by gas chromatography

After the cessation of growth, the total volume of gas produced by each culture was measured using a glass syringe, which also normalized the pressure to 1 atm. The amount of CH_4_ produced was determined by injection of 50 μl of headspace gas into a Shimazdu Nexis GC-2030 gas chromatograph fitted with a Rt-Q-BOND fused silica PLOT column with a 0.32 mm internal diameter, a 30 m length, and a 10.00 μm film thickness (Restek, VWR #89166-308) and BID detector. The sample split ratio was 42.6, and the carrier gas was helium at 4.44 mL/min. The injection port temperature was 100 °C, column temperature 27 °C, and BID temperature 220 °C. Peak integration was performed using Shimadzu LabSolutions software and moles of CH_4_ determined using methane standards.

### Data availability

The raw data from growth studies and qPCR will be available upon request.

